# Single cell reconstruction of human basal cell diversity in normal and IPF lung

**DOI:** 10.1101/2020.06.19.162305

**Authors:** Gianni Carraro, Apoorva Mulay, Changfu Yao, Takako Mizuno, Bindu Konda, Martin Petrov, Daniel Lafkas, Joe R. Arron, Cory M. Hogaboam, Peter Chen, Dianhua Jiang, Paul W. Noble, Scott H. Randell, Jonathan L. McQualter, Barry R. Stripp

## Abstract

**Rationale:** Declining lung function in patients with interstitial lung disease is accompanied by epithelial remodeling and progressive scarring of the gas-exchange region. There is a need to better understand the contribution of basal cell hyperplasia and associated mucosecretory dysfunction to the development of idiopathic pulmonary fibrosis (IPF).

**Objectives:** We sought to decipher the transcriptome of freshly isolated epithelial cells from normal and IPF lung to discern disease-dependent changes within basal stem cells.

**Methods:** Single cell RNA sequencing was used to map epithelial cell types of the normal and IPF human airway. Organoid and ALI cultures were used to investigate functional properties of basal cell subtypes.

**Measurements and Main Results:** We found that basal cells included multipotent and secretory primed subsets in control adult lung tissue. Secretory primed basal cells include an overlapping molecular signature with basal cells obtained from distal lung tissue of IPF lungs. We confirmed that NOTCH2 maintains undifferentiated basal cells and restrict basal-to-ciliated differentiation, and present evidence that NOTCH3 functions to restrain secretory differentiation.

**Conclusions:** Basal cells are dynamically regulated in disease and are specifically biased towards expansion of the secretory primed basal cell subset in idiopathic pulmonary fibrosis. Modulation of basal cell plasticity may represent a relevant target for therapeutic intervention in IPF.

## Introduction

Human airways are lined by a mucociliary epithelium that functions in the clearance of inhaled pathogens, particulates and toxic gases. Major portions of this epithelium are pseudostratified, with basal cells located adjacent to the basement membrane and subtending specialized luminal cell types including ciliated cells and various secretory cell types. Both basal and secretory cell types have proliferative capacity, but only basal cells have the capacity for long-term self renewal that is commonly associated with tissue stem cells(1, 2). Pathological remodeling of the airway epithelium is common in chronic lung disease(3, 4). Disease-related changes include hyperplasia of basal cells and altered differentiation of secretory cells leading to overproduction of mucus(5, 6).

Idiopathic pulmonary fibrosis (IPF) is a progressive lung disease with mean survival after initial diagnosis of approximately three years(7). Declining lung function results from scarring and remodeling of the interstitium surrounding the alveoli. Other pathological sequelae of IPF include the appearance of honeycomb cysts at sites of extensive parenchymal remodeling; structures that are composed of basal cells and mucin-producing airway secretory cells (5). These structural defects are considered to be secondary to epithelial progenitor cell dysfunction and defective epithelial-mesenchymal signaling (8, 9); furthermore detection of basal cells within bronchoalveolar lavage fluid of patients with IPF is associated with poor prognosis (10).

Herein, we comprehensively assess the transcriptome of single cells from normal human lung and lung tissue of patients undergoing transplantation for end-stage IPF. We found that basal cells include multiple molecularly distinct states in the normal lung, including multipotent and secretory cell primed, and present evidence suggesting that basal cell subsets form a hierarchy that is regulated by Notch signaling.

## Methods

### Study population

Explant tissue was obtained from patients undergoing transplantation for end-stage IPF at either Duke University or Cedars-Sinai Medical Center, in compliance with consent procedures accepted by the Internal Review Board of Cedars-Sinai Medical Center. Human lung specimens obtained through the International Institute for the Advancement of Medicine (IIAM) were obtained in compliance with consent procedures developed by IIAM and approved by the Cedars-Sinai Medical Center IRB.

### Data availability

All transcriptome data were deposited in GEO: GSE143706 and GSE143705. A GitHub repository with a markdown to show our QC and data analysis, is available at https://github.com/gc-github-bio/Normal-and-IPF-lung.

### Histology

Lung tissue was formalin-fixed, paraffin-embedded and sectioned at 4 μm. Cultures were fixed with 4% paraformaldehyde and washed with Phosphate-Buffered Saline (PBS) prior to whole-mount staining. Paraffin sections were deparaffinized in xylene and rehydrated through a gradient of ethanol. Prior to antibody staining, antigen retrieval was performed in 10 mM citrate buffer (pH 6.0) using a 2100-Retreiver (Aptum Biologics). Slides were washed with Tris-Buffered Saline (TBS) and blocked for 1 hour at room temperature using blocking buffer containing 3% BSA, 0.1% Triton X-100 and 10% serum (from secondary antibody species) in TBS. Primary antibodies MUC5B (HPA008246), MUC5AC (45M1, Thermofisher), FOXJ1 (14-9965-82, Biolegend), KRT5 (905904, Biolegend), KRT8 (TROMA-1, Hybrodoma Bank, Iowa), were diluted in 3% BSA, 0.1% Triton X-100 and 1% serum in TBS, applied to tissue sections and incubated overnight at 4°C. Nuclei were counterstained with 2-(4-Amidinophenyl)-1*H*-indole-6-carboxamidine (DAPI) and appropriate secondary antibodies conjugated to fluorochromes (Thermo Fisher Scientific) were applied for 1 hour at room temperature. Sections were mounted using Fluoromount G mounting medium (Thermo Fisher Scientific). Stained samples were imaged on a Zeiss LSM 780 Confocal Microscope and image processed using Zen Blue software (Zeiss).

### Cell isolation

Human IPF explants were processed to obtain epithelial cells from fibrotic regions. Donor explants were processed to obtain airway epithelial cells. The tissue was processed as described previously(1) with the following modifications. Proximal tissue was enzymatically digested with Liberase followed by gentle scraping of epithelial cells off the basement membrane. IPF tissue was finely minced and washed in Ham’s F12 (Corning) at 4°C for 5 minutes with rocking, followed by centrifugation for 5 minutes at 600g and 4°C. The minced cleaned tissue was then incubated in DMEM/F12 (Thermo Fisher Scientific) containing 1X Liberase (Sigma-Aldrich), incubated at 37°C with rocking for 45 minutes. Dissociated single-cell preparations were enriched for epithelial cells and depleted of erythrocytes, leukocytes, and endothelial cells using antibodies against the following molecules: EPCAM (CO17-1A, 369820), CD235a (HI264, 349106), CD45 (2D1,368522), and CD31 (WM59,303124) (Biolegend). For the basal cells enrichment experiment and subpopulations isolation, CD271 (ME20.4, 345106) and CD66 (ASL-32, 342306) (Biolegend) were also included. Labeled cells were washed in HBSS with 2% FBS, resuspended and placed on ice for fluorescence-activated cell sorting (FACS) using a BD Influx cell sorter (Becton Dickinson). Viability was determined by staining cell preparations with either 7AAD (Biolegend), Propidium Iodide (Biolegend) or DAPI (ThermoFisher Scientific), 15 minutes prior to cell sorting.

### In vitro cultures

Two thousand FACS enriched cells were resuspended with a 1:1 (v/v) ratio of Matrigel (Corning) and culture medium and 100uL seeded onto the apical surface of a 12 mm, hydrophilic PTFE 0.4 μm pore-size cell culture insert (EMD Millipore) placed inside a 24-well flat-bottom plate. After polymerization of the Matrigel, 500μL of basal medium (PneumaCult™-Ex Medium, STEMCELL Technologies) was added to the well, outside the cell culture insert. Cultures maintained at 37°C in a humidified incubator (5% CO_2_) and continued to be maintained in basal medium with media changes every other day. To evaluate differentiation potential and self-renewal capacity of CD66^+^, CD66^-^ basal cells, cultures were grown for up to fourteen days, at which time they were either fixed for histological analysis or organoids harvested and dissociated to yield single cell suspensions for serial passage. Colony forming efficiency was determined by counting the number of colonies with a diameter of ≥ 50μm in each culture and shown as a percentage of input epithelial cells from images taken with the EVOS^®^ XL Core transmitted-light inverted imaging system (ThermoFisher Scientific). FIJI was used to quantify colony forming efficiency from three independent experiments. Statistical analysis was performed using GraphPad Prism version 7.0a. For testing significance of the results, one-way ANOVA test was used. For serial passaging experiments, whole cultures dissociated by Liberase (Sigma-Aldrich) at 0.25 Wünsch U/mL in HBSS warmed to 37°C freed organoids from the Matrigel. After 15 minutes incubation at 37°C, cells were placed on a Thermomixer (Eppendorf) and agitated at 1000 rpm at 37°C for 20 minutes. Cells pelleted by centrifugation at 400g for 5 minutes at 4°C, followed by another dissociation step using 2mL of TrypLE™ Express Enzyme (Gibco) to generate single-cell suspensions. Once single-cell suspensions were attained and visually confirmed under light microscopy, cells were washed in HBSS-2%FBS to stop the reaction. Live cell counts were performed on resuspended cells in 0.5-1mL HBSS-2%FBS using a hemocytometer and Trypan Blue (Sigma-Aldrich). For air liquid interface (ALI) culture experiments, isolated CD66^+^ and CD66^-^ basal cells were expanded and seeded onto the apical surface of a 12 mm, hydrophilic PTFE, 0.4 μm cell culture inserts (EMD Millipore) coated with matrigel and placed inside a 24-well flat-bottom plate. Cells were grown with 500μL of basal or ALI medium (STEMCELL Technologies) according to manufacture instructions. Cultures were maintained at 37°C in a humidified incubator (5% CO_2_). For studies of NOTCH signaling, the γ-secretase inhibitor (2*S*)-*N*-[(3,5-Difluorophenyl)acetyl]-L-alanyl-2-phenyl]glycine1,1-dimethylethyl ester (DAPT) (Tocris) was added at a concentration of 5 μM. Blocking antibodies for NOTCH1 (NR1), NOTCH2 (NR2), or NOTCH3 (NR3), were added individually at concentrations of 30 μg/ml starting at the beginning of the culture, and media was changed every three days. Notch blocking antibodies were provided by Genentech.

### High-throughput FACS analysis

Samples of bronchi, bronchiolar and distal lung tissue were pooled, dissociated into a single cell suspension and used for cell screen analysis. Basal cell subset stained with APC anti-human CD271, AF488 anti-human CD326, Brilliant Violet 421 anti-human CD45 and CD31 combined with a panel of cell-surface proteins targeting 332 PE-conjugated monoclonal antibodies (Biolegend, LEGENDScreen Human PE Kit, 700007) and 10 isotype controls (Biolegend) identified novel candidate markers. Cell preparation for screening analysis were followed per manufacturer’s protocol (Biolegend). In brief, cells filtered through a 40μm cell strainer were resuspended at a density of between 1.5 – 2 x 10^6^ cells/mL in 30mLs of HBSS-2%FBS. Prior to sample staining, cells pre-incubated with human Fc-receptor blocking solution (BioLegend) to reduce nonspecific staining carried out at a concentration of (0.01%) for 15 minutes on ice. After lyophilized antibodies on plates reconstituted, plates ready for sample staining had 75μL/well of cells aliquoted and incubated in the dark for 30 minutes on ice. Cells were washed twice and fixed for 10 minutes at room temperature prior to flow cytometer analysis. Analyses were performed on labeled cells using an LSRFortessa cell analyzer (Becton Dickinson, BD) and collected data collected were interrogated using FlowJo software.

### Population RNAseq

Basal cell CD66^+^ and CD66^-^ populations were processed immediately or after in vitro culturing for ten days. RNA was extracted from the sorted cells using the RNeasy Micro Kit (Quiagen). Total RNA was quantified and sample contamination by proteins or carryover reagents were assessed using both a NanoDrop and a Qubit (Thermo Scientific). Samples were then qualified using the Fragment Analyzer (Advanced Analytical Technologies) to check the integrity of total RNA by measuring the ratio between the 18S and 28S ribosomal peaks. SMART-Seq V4 Ultra Low RNA Input Kit for Sequencing (Takara Bio USA, Inc., Mountain View, CA) was used for reverse transcription and generation of double stranded cDNA for subsequent library preparation using the Nextera XT Library Preparation kit (Illumina, San Diego, CA). Reaction Buffer (1X) was added to lysed cells and the RNA was used directly for oligo(dT)-primed reverse transcription, followed by cDNA amplification and cleanup. Quantitation of cDNA was performed using Qubit (Thermo Fisher Scientific). cDNA normalized to 80 pg/μl was fragmented and sequencing primers added simultaneously. A limiting-cycle PCR added Index 1 (i7) adapters, Index 2 (i5) adapters, and sequences required for cluster formation on the sequencing flow cell. Indexed libraries were pooled and cleaned up, and the pooled library size was verified on the 2100 Bioanalyzer (Agilent Technologies) and quantified via Qubit. Libraries were sequenced on a NextSeq 500 (Illumina) using with a 1×75 bp read length and coverage of over 25M reads per sample. Population RNAseq reads were processed using packages inside Galaxy (https://usegalaxy.org/). Specifically, alignment to human genome hg38 was performed using STAR 2.6.0b-1 with application of the default setting in Galaxy. Raw counts were measured using FeatureCounts 1.6.0.3 with application of the default setting in Galaxy. The annotation GTF file of USCS hg38 was downloaded from iGenomes (https://support.illumina.com/sequencing/sequencing_software/igenome.html).

Differential gene expression was determined using DESeq2(4) (version 2.11.40.2) using default parameters in Galaxy. The Top DEG for freshly isolated and cultured datasets, along with raw and normalized counts, are shown in Table 6.

Heatmaps in Fig. 3C-D were produced selecting the top 20 differentially expressed genes in each group on the base of their log2 fold change among genes with adjusted p-values < 0.05. Heatmaps in Fig. S4A-B were produced as follows: for popRNAseq DEG, only genes with adjusted p-value < 0.05 were used for analysis. A DEG of scRNAseq data of MPB and SPB cells was produced (Table 6) and only genes with adjusted p-value < 0.05 were used in the analysis. scRNAseq DEG were used to define PopRNAseq DEG that were used in the analysis. For freshly isolated cells the top 20 DEG’s for CD66+ and CD66-where used for creation of the heatmap. For popRNAseq of cultured basal cells, only 13 genes passed the filtering and were used for creation of the heatmap.

### Single cell RNAseq

Single cells were captured using a 10X Chromium device (10X Genomics) and libraries were prepared according to the Single Cell 3’ v2 or v3 Reagent Kits User Guide (10X Genomics, Table 9 specifies the kit version used for each dataset). Totalseq-A human hashing antibodies (Biolegend) were used to label cells isolated from sample cc05; trachea or extralobar bronchi used TotalSeq-A0253 (394605), proximal intra-lobar bronchi used TotalSeq-A0254 (394607). Hashing was performed for a different purpose and was not considered for data processing in this manuscript. Cellular suspensions were loaded on a Chromium Controller instrument (10X Genomics) to generate single-cell Gel Bead-In-EMulsions (GEMs). Reverse transcription (RT) was performed in a Veriti 96-well thermal cycler (ThermoFisher). After RT, GEMs were harvested, and the cDNA underwent size selection with SPRIselect Reagent Kit (Beckman Coulter). Indexed sequencing libraries were constructed using the Chromium Single-Cell 3’ Library Kit (10X Genomics) for enzymatic fragmentation, end-repair, A-tailing, adapter ligation, ligation cleanup, sample index PCR, and PCR cleanup. The barcoded sequencing libraries were quantified by quantitative PCR using the KAPA Library Quantification Kit (KAPA Biosystems, Wilmington, MA). Sequencing libraries were loaded on a NovaSeq 6000 (Illumina) with a sequencing setting of 26bp and 98bp for v2 and 28bp and 91bp for v3, respectively, to obtain a sequencing depth of at least ~5 x 10^4^ reads per cell.

### Data analysis

Cell Ranger software (10X Genomics) was used for mapping and barcode filtering. Briefly, the raw reads were aligned to the transcriptome using STAR(5), using a hg38 transcriptome reference from Ensembl 93 annotation. Expression counts for each gene in all samples were collapsed and normalized to unique molecular identifier (UMI) counts. The result is a large digital expression matrix with cell barcodes as rows and gene identities as columns.

Data analysis was mainly performed with Seurat 3.0(6) with some variation that will be described.

For all data, quality control and filtering were performed separately for each dataset, to remove cells with low number of expressed genes (threshold n>=200) and elevated expression of apoptotic transcripts (threshold mitochondrial genes < 15%). Only genes detected in at least 3 cells were included. Each dataset was run separately with SoupX analysis package to remove contaminant ‘ambient’ RNA derived from lysed cells during isolation and capture (Young MD et al., https://doi.org/10.1101/303727). We performed a correction on the basis of genes with a strong bimodal distribution and for which the ‘ambient’ RNA expression was overlapping with a gene signature of a known cell type. The ‘adjustCounts’ function of SoupX was used to generate corrected count’s matrices. Table 8 show the percentage of ambient RNA identified in each dataset. To minimize doublet contamination for each dataset we performed a quantile thresholding of high UMI using a fit model generated using the multiplet’s rate to recovered cells proportion, as indicated by 10X Genomics (https://kb.10xgenomics.com/hc/en-us/articles/360001378811-What-is-the-maximum-number-of-cells-that-can-be-profiled-). The raw expression matrix was processed with SCTransform wrapper in Seurat. Each dataset was first processed separately with Principal Component Analysis (PCA), followed by clustering using the first 30 independent components and a resolution of 0.5. Two-dimensional visualization was obtained with Uniform Manifold Approximation and Projection (UMAP)(8). Identified AT2 (SFTPC+), immune (PTPRC+), and endothelial (PECAM1+) contaminating clusters were removed by subsetting the Seurat object, using the ‘subset’ function, before proceeding to merging of the data and integration. In the merged raw expression matrix, each sample was processed with SCTransform. Mitochondrial and ribosomal mapping percentages were regressed to remove them as source of variation. We obtained log1p logarithmically transformed data for each dataset and scaled data as Pearson residuals. Pearson residual was then used to integrate datasets following Seurat workflow, using the PrepSCTIntegration function. Integrated datasets were used for downstream analysis. The integrated datasets didn’t show batch effects from sample of origin allowing datasets clustering on the base of biological significance (Fig. S1F-G). Datasets were processed with PCA using the 5000 most variable genes as input, followed by clustering with Leiden algorithm using the first 30 independent components and a resolution of 3 for fine clustering. Two-dimensional visualization was obtained with UMAP. To identify differentially expressed genes between clusters we used Model-based Analysis of Single-cell Transcriptomics (MAST)(9) within Seurat’s FindMarkers and FindAllMarkers functions. For this analysis the p-value adjustment is performed using Bonferroni correction based on the total number of genes. Results for major clusters and subclusters are reported as supplementary tables (Supplementary Table 2, 3). To identify major cell types in our normal integrated datasets, we used previously published lung epithelial cell type specific gene lists(10) (Supplementary Table 1) and created a scoring, using a strategy previously described(11). This method calculates the expression of each gene signature list as average, and this is subtracted by the aggregated expression of control feature sets. All analyzed features are binned based on averaged expression, and the control features are randomly selected from each bin. Thirty-seven clusters identified with the Leiden algorithm were assigned to major cell types on the base of rounds of scoring and refinement (Fig S2). Each refinement was produced using transcripts differentially expressed within the best identified clusters from the previous scoring. Within each major cell type, Leiden clustering and differential gene expression were used to infer subclustering. Subclustering segregation was visualized by heatmap showing the top differentially expressed genes, and by violin plot, showing a signature score obtained as described above (Fig 1). The gene list used as signature are shown in supplementary tables (Supplementary Table 2, 3). Violin plots show expression distribution and contain a boxplot showing median, interquartile range, and lower and upper adjacent values. To identify CD markers that can discriminate basal cell subtypes, we checked all CD markers (from HGNC, https://www.genenames.org) (Supplementary Table 4) with differential expression in our datasets, and selected markers that were confirmed by FACS screening.

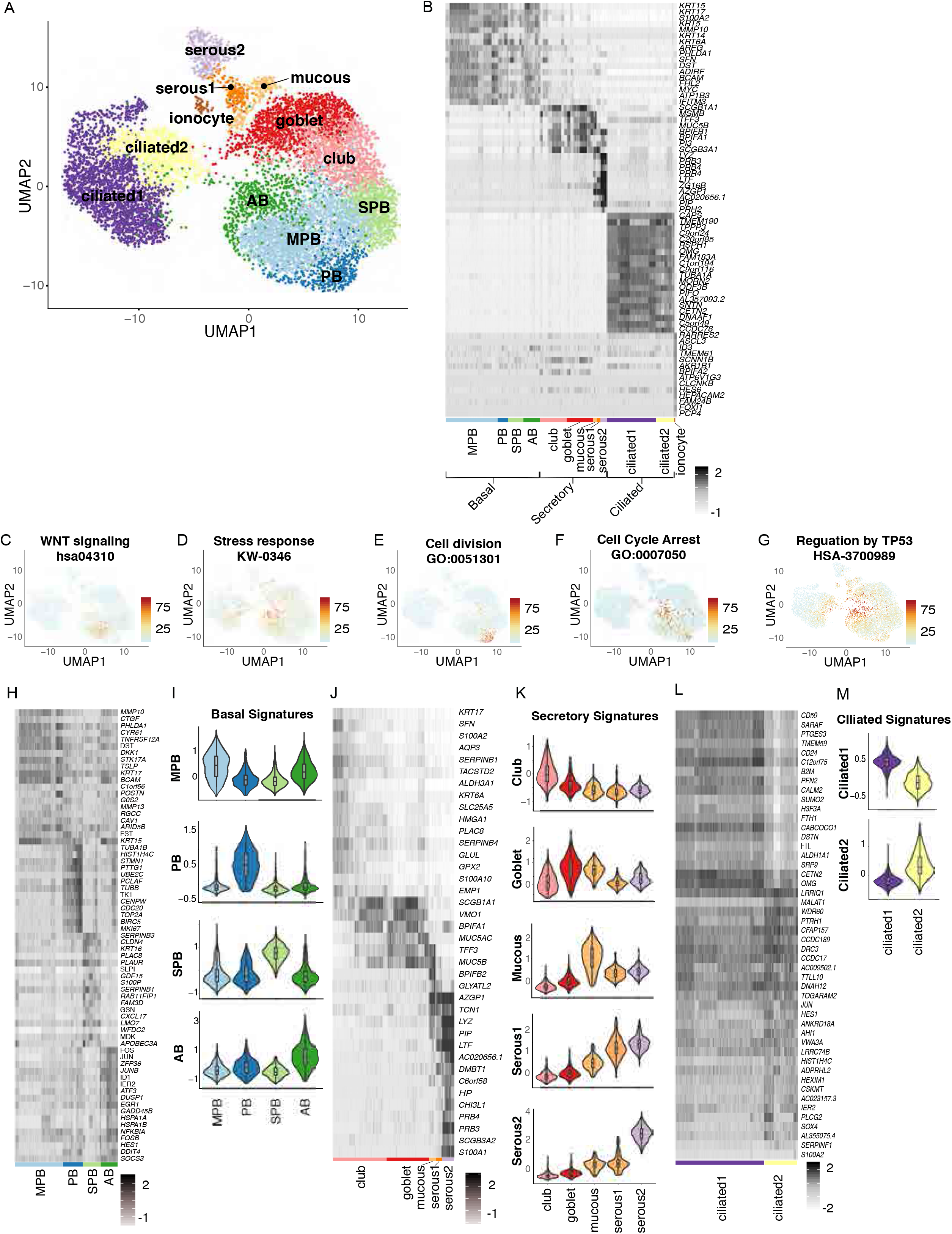

Top differentially expressed genes were fed to STRING (https://string-db.org/) to visualize their protein association network and extract their functional enrichments. Gene list of specific terms were used to create a score, as described above, and visualized as UMAP. IPF datasets were first processed as described for normal datasets, from QC analysis to integration. For comparative analysis of control and IPF, a common integration of all datasets was performed following the same integration strategy described for control datasets. The integrated Seurat object containing control and IPF data was used to produce differential gene expression analysis using MAST (Supplementary Table 7). This analysis was used to generate visualizations in Fig.4 B and G.

To estimate the presence of cellular transitions between basal and secretory clusters, we used diffusion map with ‘Destiny’(12), followed by trajectory analysis with ‘Slingshot’(13), following their Bioconductor vignettes. The normalized counts from Seurat were imported into an ExpressionSet for Destiny and into a SingleCellExperiment for Slingshot. The 3D visualization was obtained using the ‘rgl’ library in R. Notch signaling ligand-receptor interactions were analyzed using iTALK (Wang Y et al., https://doi.org/10.1101/507871). All analyses were performed using R 3.6.

### Statistical analysis

FIJI was used to quantify colony forming efficiency from three biological samples in at least eight technical replicates. Statistical analysis was performed using GraphPad Prism version 7.0a. Data presented are shown as mean ratio ± SEM. For testing significance of the results, one-way ANOVA test was used. For quantification of immunofluorescence experiments with at least three biological replicates Mann-Whitney test (P<0.1) was used.

For analysis of single-cell transcriptomics the p-value adjustment was performed using Bonferroni correction based on the total number of genes. For population RNAseq all samples were collected and stored to allow processing at the same time. For single-cell RNAseq, samples were processed based on sample availability, so balanced batch preparation could not be performed.

## Author contributions

G.C. and B.R.S. designed the project and wrote the manuscript. G.C. performed scRNAseq data analysis. A.M. designed and performed experiments for Notch signaling. T.M. designed and performed organoid experiments and population RNAseq. M.P. and B.K. performed tissue processing and immunostainings. C.Y. performed single-cell RNAseq experiments and popRNAseq data processing, M.P. performed single-cell RNAseq experiments. J.L.M. performed high throuput FACS screening experiments and contributed to the project design. D.L., J.R.A. and S.R. contributed to data interpretation. P.C., D.J., and P.W.N. contributed with comments on the manuscript. All authors read and reviewed the manuscript.

## Results

### Mapping the molecular phenotype of normal human lung airway epithelial cells

Single-cell RNA-sequencing (scRNA-seq) was performed on fractionated epithelial cells from six donors with no preexisting chronic lung disease and no or mild smoking history (Fig S1A). A single-cell transcriptome of the aggregated datasets containing 9731 cells was generated after initial filtering. We defined 37 clusters (Fig S2A) encompassing 12 cellular phenotypes, including 4 basal, 5 secretory, 2 cilated, and ionocyte cell types, that were visualized by Uniform Manifold Approximation and Projection (UMAP) and by their top differentially expressed genes (Fig 1A, B, Table 2). Clusters were first assigned to major cell populations using previously published gene lists(11) (Table 1) to create a signature score(12). Clustering and subclustering were than refined by multiple iterations of scoring, based on differential gene expression (Fig S2C-L). Clusters 18, 28, and 31, initially uncharacterized, were recognized as secretory cell types after the first round of score refinement (Fig S2D), and were than manually annotated as mucous and serous cells on the base of published markers for submucosal gland(13) (Fig 1A, J, K). Using the same approach two separate clusters of ciliated cells were identified. A relatively immature (ciliated2) cluster that contains genes of early ciliogenesysis such as *LRRC6, DNAAF1,* and *DNAAF5* (Fig 1A, L, M, FigS1 J) and a mature ciliated1 cluster (Fig 1A, L, M). Each cell type classification was represented by single cells from each donor (Fig S1B).

### Normal human airway basal cells comprise specialized subpopulations

Basal cells segregated into 4 clusters on the basis of their gene signatures (Fig 1A, B, H, I; Table 3). MPB cluster represents a ‘classical’ Multipotent Basal (MPB) cell that was defined by a strong WNT signaling signature (Fig 1A-C). PB cluster represents proliferating basal (PB) cells (Fig 1A, B, E). Cells within the SPB cluster express various members of the serpin family (Fig 1H, S1I) that regulate protein folding in the early secretory pathway(14). SPB cells also show enrichment of transcripts involved in O-glycan biosynthesis (Fig 2A), a pathway involved in the initial steps of mucin production by secretory cells (15). The Activated Basal (AB) cluster is characterized by a stress-response signature (Fig 1A, B, D) including p38-SAPK MAP kinase cellular stress (Fig S1K). SAPK has been involved in centriolar satellite remodeling and ciliogenesis (16). Furthermore AB cluster is also characterized by activation of the TP53 pathway (Fig 1G) and contain a cell-cycle arrest signature (Fig 1F).

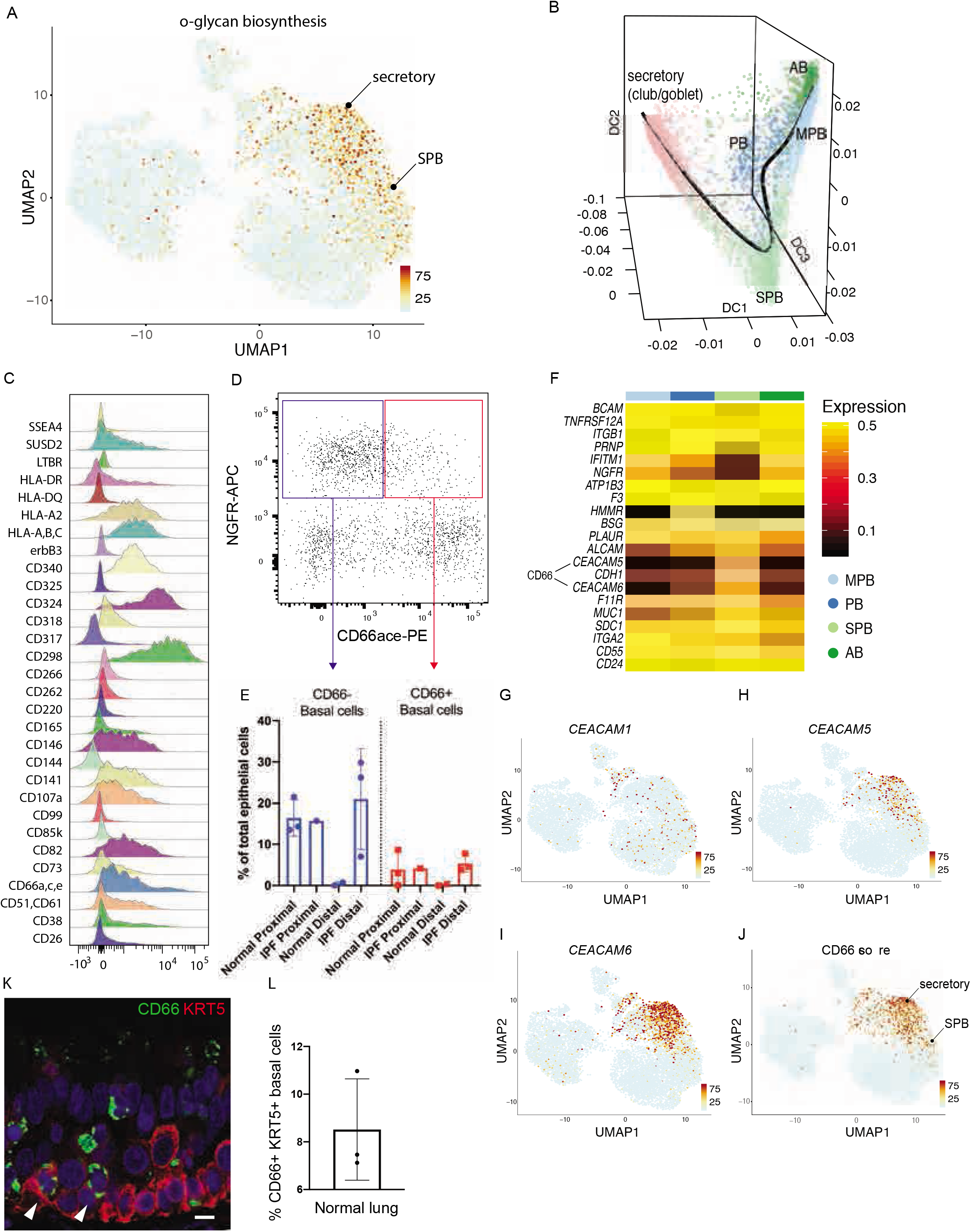

In addition to scRNAseq we used a high content flow cytometry screening to detect basal cell heterogeneity based on surface marker expression. This discovery screen was performed on pooled samples of epithelial cells isolated from bronchi, bronchiolar and distal lung regions from a single IPF patient. The requirement for very high cell numbers for this type of high throughput screen restricted our capacity to perform this screen on separate spatially localised samples. A total of 332 individual surface markers (plus isotype controls) were screened in combination with a viability die and antibodies against CD31 (endothelial cells), CD45 (hematopoietic cells), EPCAM (epithelial cells) and NGFR (basal cells). Analysis for distribution of cells in the CD31-CD45-EpCAM+ NGFR+ basal cell fraction revealed 30 surface markers that could be used for basal cell subclustering (Fig 2C, S3). Of these, CD66 recognition of CEACAM1,5,6 isoforms was one of the markers that was identified to correlate with selective expression of *CEACAM 5* and *6* by the SPB cluster in the scRNAseq data (Fig 2F) and was thus chosen for further validation in normal (n=3) and IPF (n=3) lungs (Fig 2D, E). This confirmed differential surface reactivity of CD66 among basal cells in the normal proximal lung, and showed a trend for an elevated proportion of CD66+ basal cells in IPF. Immunofluorescent staining of normal lung airways showed that CD66 immunoreactivity was observed at the plasma membrane in a rare population of basal cells in addition to previously described subpopulations of luminal transitional and secretory cells (Fig 2K, L, S4C) (17). Expression of *CEACAM 5* and *6* isoforms that contribute to CD66 immunoreactivity of SPB and secretory cells is also observable at the transcriptional level by UMAP (Fig 2G-J). Interestingly, dimensional reduction by diffusion map with ‘Destiny’ and lineage visualization with ‘Slingshot’ show that SPB cells have a molecular phenotype that is transitional between basal and secretory cells (Fig 2B). However, our analysis does not allow determination of whether this population represents a transitory or phenotypically stable subset of basal cells.

### SPB cells have limited capacity for self-renewal and are primed to assume secretory cell fates

We next sought to test the functional potential of SPB cells and validate computational predictions from scRNAseq data. Total basal cells were sorted for either EPCAM^+^, NGFR ^+^, CD66^+^ (CD66^+^ cells) or EPCAM^+^, NGFR^+^, CD66^-^ (referred to as CD66^-^ cells). Mesenchyme-free three-dimensional in vitro cultures were used to assess the capacity of basal cell subsets to form clonal organoids and evaluated differentiation potential. Both CD66^+^ and CD66^-^ populations generated clonal organoids in the absence of mesenchymal support. However, CD66^+^ organoids displayed reduced capacity for selfrenewal starting at passage 2, compared to their CD66^-^ counterparts (Fig. 3A). We performed bulk RNA-seq on either freshly isolated or cultured basal cell fractions. Top differentially expressed genes (log2 fold change, adjusted p-value < 0.05) for CD66^+^ and CD66^-^ are shown (Fig. 3C, D, Table 6). Top differentially expressed genes (log2 fold change, adjusted p-value < 0.05) that were also differentially expressed in SPB and MPB cells (scRNAseq data), are shown (Fig. S4A,B, Table 6). PCA analysis showed that CD66^+^ and CD66^-^ cells diverged significantly seven days after organoid culture reflecting their disparate fates (Fig 3B). Immunofluorescence showed marked up-regulation of secretory cell markers MUC5AC and MUC5B (Fig. 3E, G, S5). When seeded at high density on Transwell^®^ cultures and grown at air-liquid interface CD66^+^ basal cells also demonstrated a more pronounced propensity for secretory cell differentiation compared to their CD66^-^ counterparts (Fig 3F, H). These data are consistent with the notion that SPB CD66^+^ basal cells are primed to assume secretory cell fates.

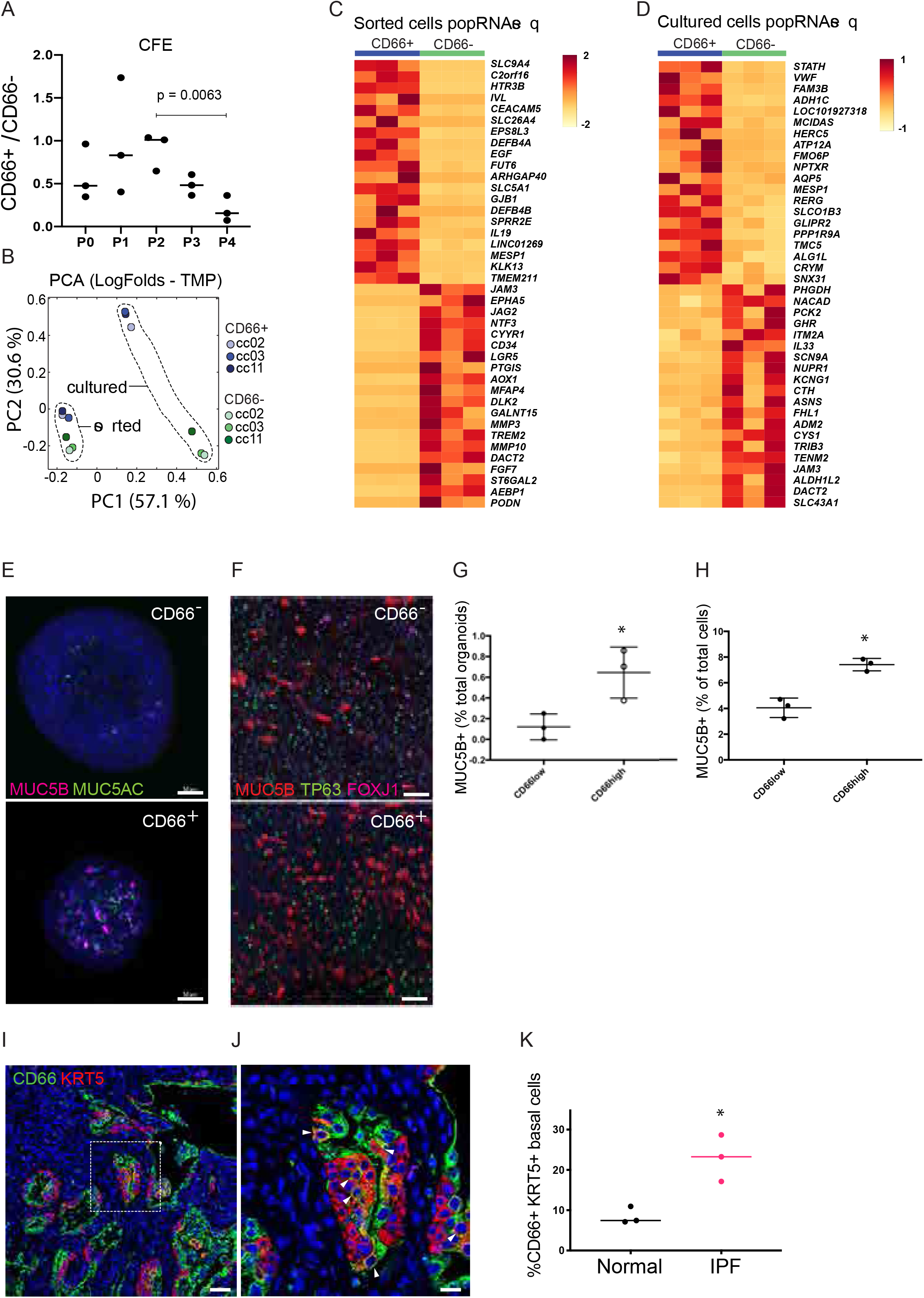

### IPF distal lung contains an expanded pool of SPB cells

We previously showed that decreases in the abundance of alveolar type 2 cells (AEC2) within explant tissue of patients undergoing transplant for end-stage IPF was accompanied by a corresponding increase in the abundance of NGFR^+^ basal cells(18). These findings were of particularl interest in light of GWAS data linking MUC5B promoter polymorphism and susceptibility to IPF, which suggest potential roles for altered airway secretory cells in disease progression(19). Immunofluorescent staining to localize CD66^+^ KRT5^+^ basal cells in end-stage IPF explant tissue revealed an increase in SPB cells (Fig 3I-K). To better define changes to basal cells in the IPF lung we evaluated scRNAseq data from IPF distal lung epithelium. Single cell transcriptomes of 7 IPF samples (Fig S1A, C, G) were generated and the initial quality control and filtering were performed as for control subjects. IPF datasets were classified by the gene signatures obtained for each of the major epithelial cell types defined in the normal lung (Fig 4A). Shared transcripts between IPF and control datasets were evaluated as integrated data, showing similarity among major cell types (Fig 4B). IPF datasets contained representative cells for AB, MPB, and SPB cells identified in the normal lung. Control and IPF cells showed an incomplete signature overlap. A specific proliferative cluster of IPF basal cells was not identified. Instead, proliferating cells were present in all IPF datasets (Fig S1H). Furthermore, two secretory cell clusters specifically enriched in IPF were named on the basis of their specificity for selected secretoglobins: SCGB3A2+ and SCGB3A2+/SCGB3A1+ (Fig. 4A). IPF datasets were enriched in CD66 expression, in SPB and in all the secretory clusters (Fig. 4 C-F, Table 7). IPF SPB cells were specifically enriched in *CEACAM6* expression (Fig 4E), and could be distinguisged from normal SPB cell by their gene signature (Fig 4G, Table 7). Dimensional reduction by diffusion map with ‘Destiny’ and lineage visualization with ‘Slingshot’ show that also in IPF datasets basal and secretory cells form a continuum, with SPB cells being the connection to secretory cells (Fig 4H).

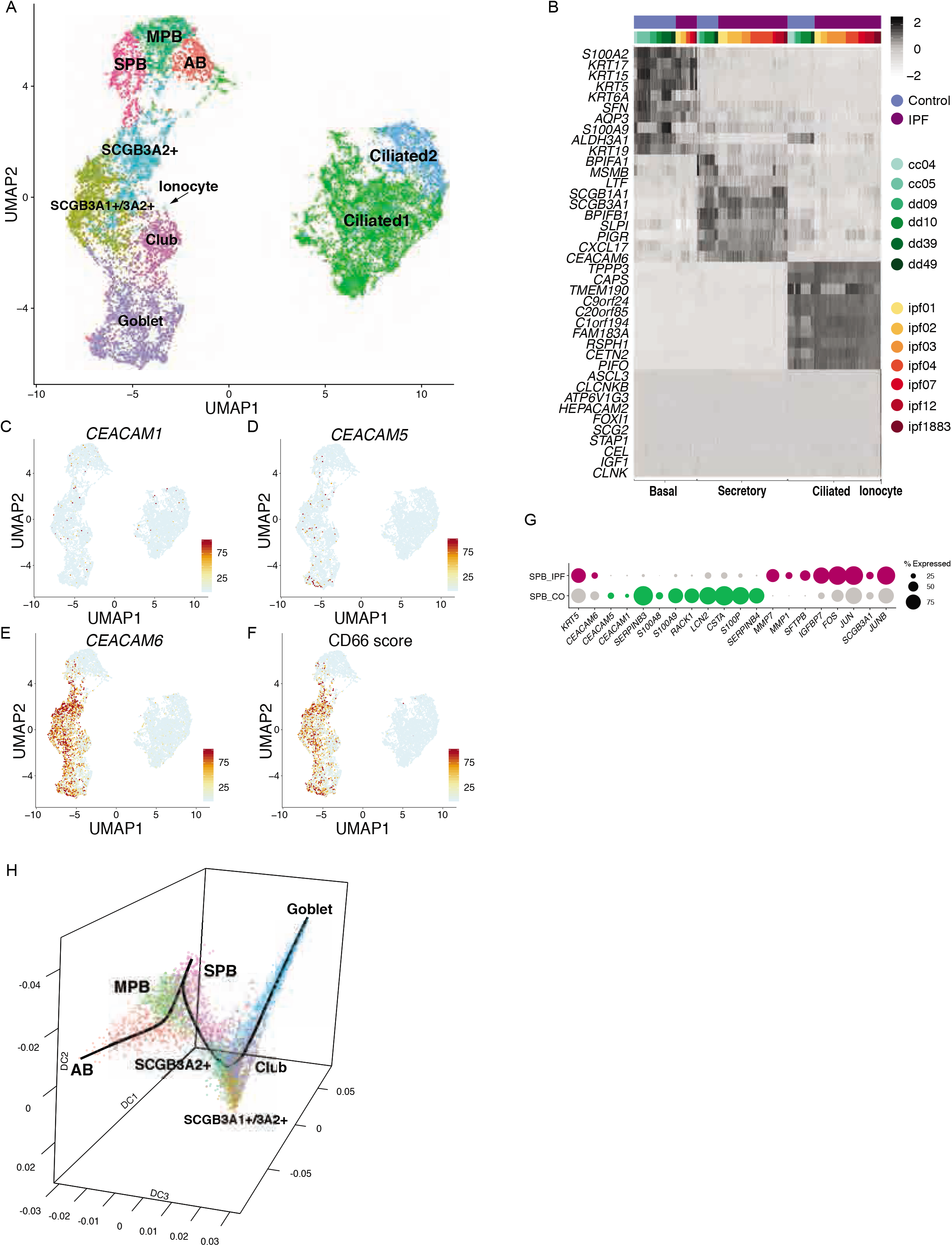

### Notch signaling regulates the maintenance of SPB cells

Given the importance of Notch signaling in regulating lung airway maintenance and differentiation, we interrogated Notch pathway gene expression between basal cell subsets of control lung. Analysis of Notch ligand-receptor interactions suggest that both signaling and receiving cells were present within the total basal cell population (Fig 5). Interestingly the SPB cluster posses the largest amount of both autocrine and juxtacrine Notch signaling (Fig 5A). *NOTCH1* was the most broadly involved receptor and was predicted to play a significant role in the regulation of all basal cell subsets with the exception of MPB cells (Fig 5B). In contrast, signaling by *NOTCH3* was the most restricted, showing significant regulation of only the SPB subset (Fig 5B, R-T). The PB cells (proliferating basal cells) showed the least signaling activity and were defined predominantly as Notch receving cells.

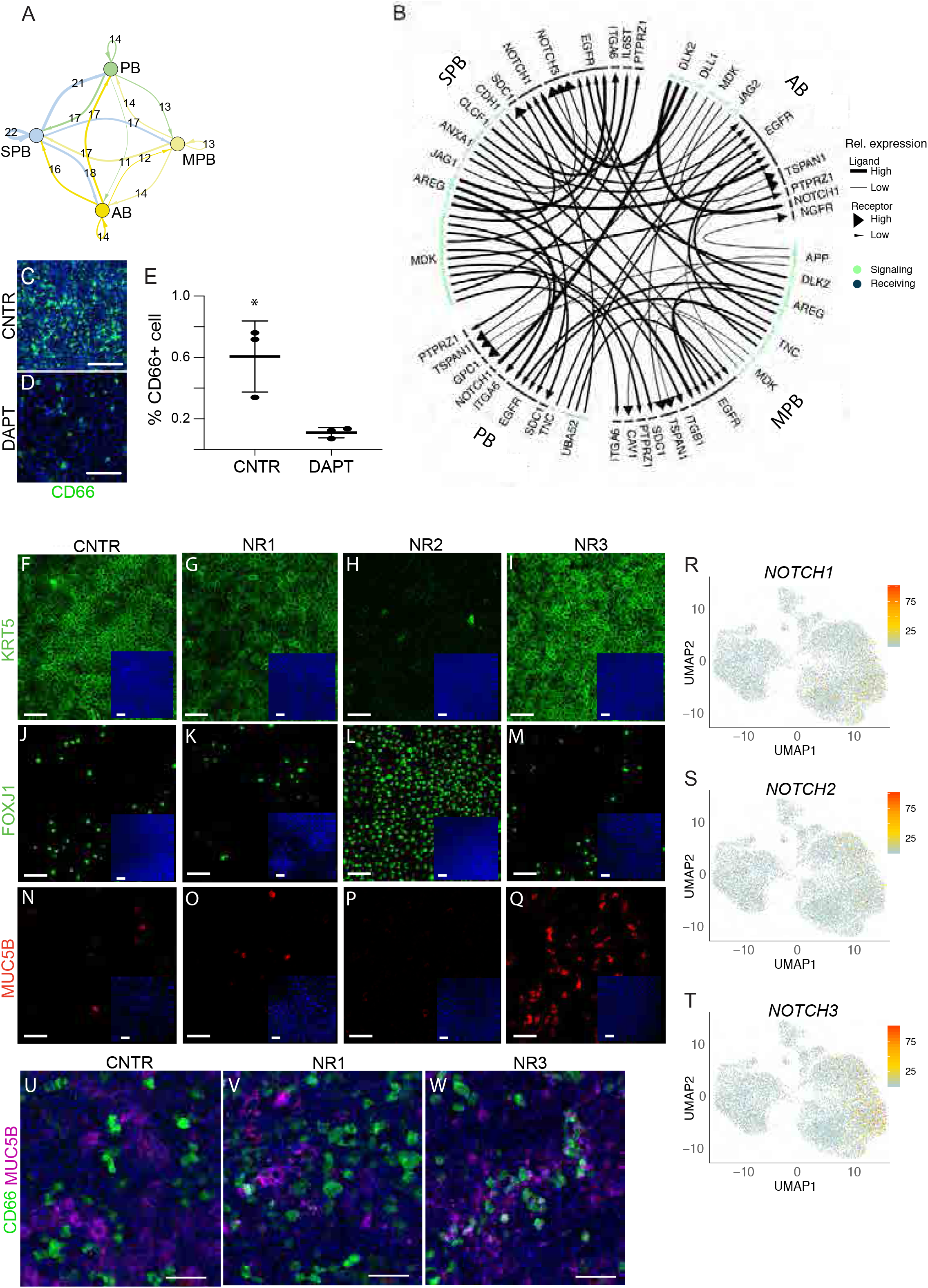

To test the importance of Notch signaling in SPB cells maintenance we established an ALI culture with a 1:1 proportion of CD66^+^ (SPB) and CD66^-^ cells. Treatment with DAPT, a γ-secretase inhibitor that blocks release of the NOTCH intracellular domain, dramatically reduced the abundance of CD66^+^ SPB cells 48h after treatment, suggesting a functional role for Notch signaling in the maintenance of SPB cells (Fig 5C-E). To specifically interrogate the significance of the three Notch signaling receptors for maintenance and differentiation of basal cells, the fate of basal cells in primary cultures of human tracheal epithelium was evaluated following treatment with specific blocking antibodies for NOTCH1 (NR1), NOTCH2 (NR2), and NOTCH3 (NR3). Cells grown in ALI culture were harvested after 24h, 8 days, and 14 days of treatment. From day 8 to day 14, NR2 treated ALI culture showed a dramatic reduction in KRT5^+^ cells compared to control (Fig 5F-I). Interestingly, neither NR1 nor NR3 treatment impacted the abundance of KRT5^+^ cells, suggesting that NOTCH2 is the main ligand involved in basal cell maintenance. Furthermore, NOTCH2 inhibition also promoted expansion of ciliated cell differentiation as shown by an increase in FOXJ1-immunoreactive cells (Fig 5J-M). These data are consistent with the previously reported effects γ-secretase inhibitors, which promote commitment of secretory progenitors to multiciliated cell fates(20) and suggest that NOTCH2 is the relevant receptor that mediates these effects. Even though inhibition of NOTCH3 had no impact on the abundance of KRT5^+^ basal or FOXJ1^+^ ciliated cells in culture, NR3 treatment led to an increase in the proportion of mucin secreting cells at 8 days (Fig 5N-Q) and 14 days (Fig 5U-W) in culture. These data suggest that NOTCH3 may specifically restrains the differentiation of SPB cells, while NOTCH2 plays a more global role in regulating basal cell maintenance and restrains ciliated cell differentiation.

## Discussion

Airway basal cells function as stem cells that maintain the pseudostratified epithelium of the mouse and presumably also the human lung. Even though basal cell heterogeneity has been implied from studies of the pseudostratified epithelium lining the mouse trachea, little is known of basal cell heterogeneity in the normal human lung and how this changes in the setting of pathological tissue remodeling that accompanies disease. Employing single cell transcriptomics of freshly isolated human airway epithelial cells we were able to discern four broad classes of epithelial cells that included basal, secretory, ciliated, and ionocyte. Even though heterogeneity was observed within each of the abundant epithelial cell types, significant heterogeneity was observed among basal cells. Basal cell subsets included a Wnt signaling multipotent cluster, proliferating basal cells, a cluster that showed evidence of secretory priming, and a cluster with activation of stress response genes. Secretory Primed Basal (SPB) cells may either represent a transitory state of basal cells that are in the process of secretory cell differentiation or a phenotypically stable subset. Diffusion mapping pseudotime analysis of differentiation trajectory show a continuum of differentiation, suggesting a transitional state. However, immunofluorescence analysis reveals rare basal cells that are positive for the SPB marker CD66 in control adult lung tissue and we also found that these cells can be clonally expanded in vitro for multiple passages in the absence of mesenchymal support. Further work is therefore necessary to exaustively determine the features of these cells and their contribution to homeostasis and disease. A recent study performing single-cell transcriptomic analysis of epithelial cells isolated from air-liquid interface cultures of nasal epithelium(21) described a KRT5+ cell type, termed ‘suprabasal’, that may represent a precursor of secretory cells. Suprabasal cells were characterized by expression of *KRT13,* a transcript observed in SPB cells in our study. Furthermore, suprabasal cells share additional similarity with SPB cells in their expression of *NOTCH3.* However, differences between transcriptomes of suprabasal and SPB cells were observed, such as expression of *KRT4* in suprabasal cells but absence of this cytokeratin in SPB cells.

These may represent cell type differences between epithelia of the upper and lower respiratory tracts or represent differences in cell states associated with isolation from fresh tissue versus cultures.

Studies in mice have established the existence of subsets of quiescent basal cells in the pseudostratified epithelium that lines the mouse trachea(22). Notch2-ICD expression was shown to define a subpopulation of basal cells that preferentially generate luminal secretory cells following injury. In contrast, c-Myb expression defined basal cells that were primed to undergo multiciliogenesis after injury(23). Our analysis of basal cells in the human lung suggests conservation of Notch signaling as a regulator of basal cell fate. In cultures of freshly isolated human basal cells we found that either γ-secretase inhibitors or isoform-specific Notch blocking antibodies biased basal cell differentiation towards secretory or ciliated phenotypes. We specifically found that NOTCH3 restrains secretory primed basal cells from assuming mature secretory cell fates and that is a global regulator of basal cell maintenance. These findings are consistent with studies in mice showing that NOTCH3 signaling regulates the pool of epithelial progenitors that are competent to respond to NOTCH receptors 1 and 2 (24). In basal cells of the human airway NOTCH2 is required for their differentiation (25). This is consistent with our observation that blocking NOTCH2 with anti-NOTCH2 antibody blocks secretory and enhances ciliated cell differentiation. Our results with anti-NOTCH3 differ from those by Danahay et al(23) in that we see an increase in secretory cell differentiation, an effect that was not observed in the Danahay study. One explanation for this discrepancy could be that we co-culture CD66+ and CD66-basal cells, hence increasing the proportion of CD66+ basal cells in culture and allowing more precise evaluation of this basal cell subset. JAG blocking antibodies were reported to produce loss of club cells and a gain in ciliated cells(26). This result is consistent with our observed effect of anti-NOTCH2 on ciliated cell expansion. We provide evidence that basal cells in fibrotic explant lung tissue obtained from patients undergoing transplant for end-stage IPF include secretory primed basal (SPB) cells that are found in distal airways and honeycomb regions of IPF lungs that are the site of excessive MUC5B expression(5). Even though the pathophysiological significance of honeycombing and excessive mucus secretion is not fully understood, Seibold et al showed that a polymorphism of the MUC5B gene is associated with increased susceptibility to IPF but surprisingly IPF patients that carry this polymorphism show better outcomes compared to those with the wildtype MUC5B allele(19). We found that IPF SPB cells express genes previously identified as serum biomarkers for higher risk of mortality(27) or proposed to be involved in disease development(28), such as *MUC1, MMP7,* and *ICAM1.*

This work defines a basal cell hierarchy that is dynamically regulated between health and disease, presenting new therapeutic targets for modulation of normal tissue maintenance and remodeling.

## Acknowledgments

We would like to thank Adrianne Kurkciyan and Stephen Beil for technical assistance, Dr. Edo Israely for assistance with FACS and Cedars-Sinai Medical Center Flow Cytometry, Genomics cores, and Biobank. Dr. Allon M. Klein and Caleb Weinreb for helpful discussions.This work was supported by grants from the California Institute for Regenerative Medicine (LA1-06915), the Cystic Fibrosis Foundation (CARRAR19G0, STRIPP15XX0), Celgene, the NHLBI (R01 HL135163, HL138540, P01 HL108793, T32 HL134637 and NIDDK (P30 DK065988).

