## Supplemental Figures for "Single cell reconstruction of human basal cell diversity in normal and IPF lung"

### Figure S1

A

| Platform | 10X Kit | Type | ID | Age | Gender | Smoking History |
| --- | --- | --- | --- | --- | --- | --- |
| 10X | v3 | Normal | cc04 | 47 | M | smoker (½ py) |
| 10X | v3 | Normal | cc05 | 63 | F | smoker (½ py) |
| 10X | v2 | Normal | dd09 | 53 | F | smoker (4.5 py) |
| 10X | v3 | Normal | dd10 | 24 | M | non-smoker |
| 10X | v3 | Normal | dd39 | 52 | M | non-smoker |
| 10X | v3 | Normal | dd49 | 48 | F | smoker (13 py) |
| 10X | v2 | IPF | ipf1883 | 64 | F | non-smoker |
| 10X | v2 | IPF | ipf01 | 64 | F | non-smoker |
| 10X | v2 | IPF | ipf02 | 62 | F | smoker (9 py) |
| 10X | v3 | IPF | ipf03 | 72 | M | smoker (quit 1/1/15) |
| 10X | v2 | IPF | ipf04 | 65 | F | non-smoker |
| 10X | v2 | IPF | ipf07 | 68 | F | non-smoker |
| 10X | v3 | IPF | ipf12 | 64 | M | non-smoker |

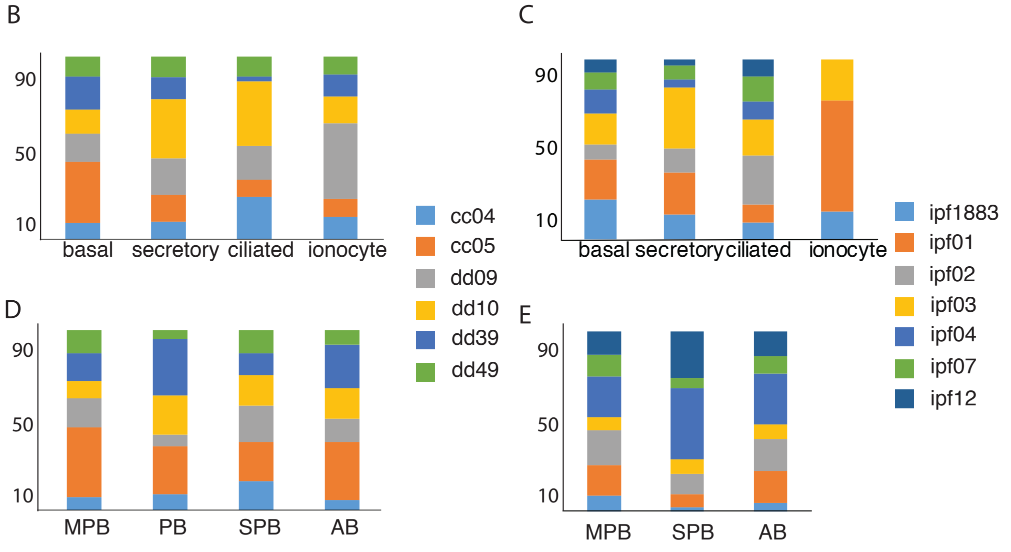


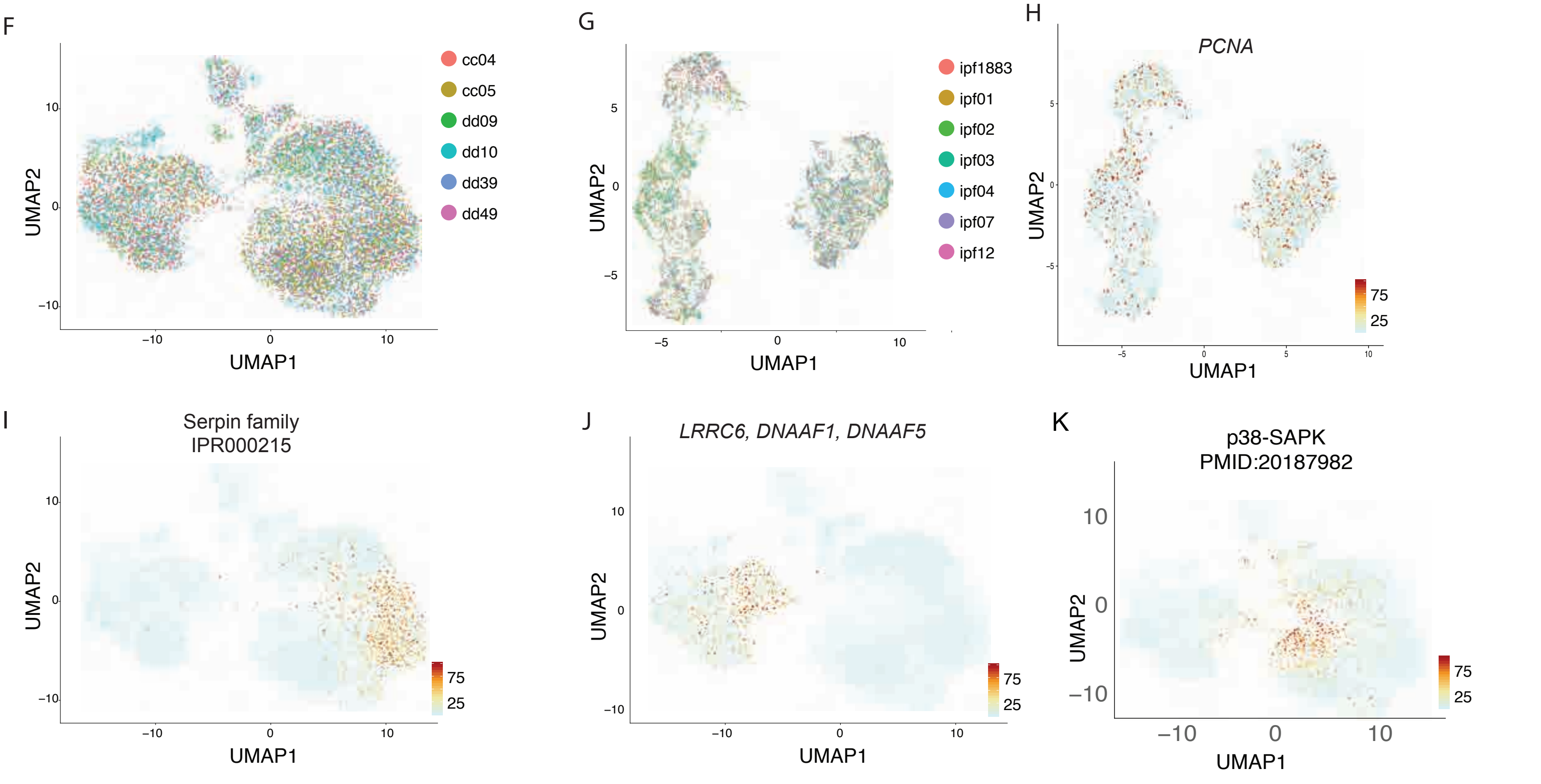

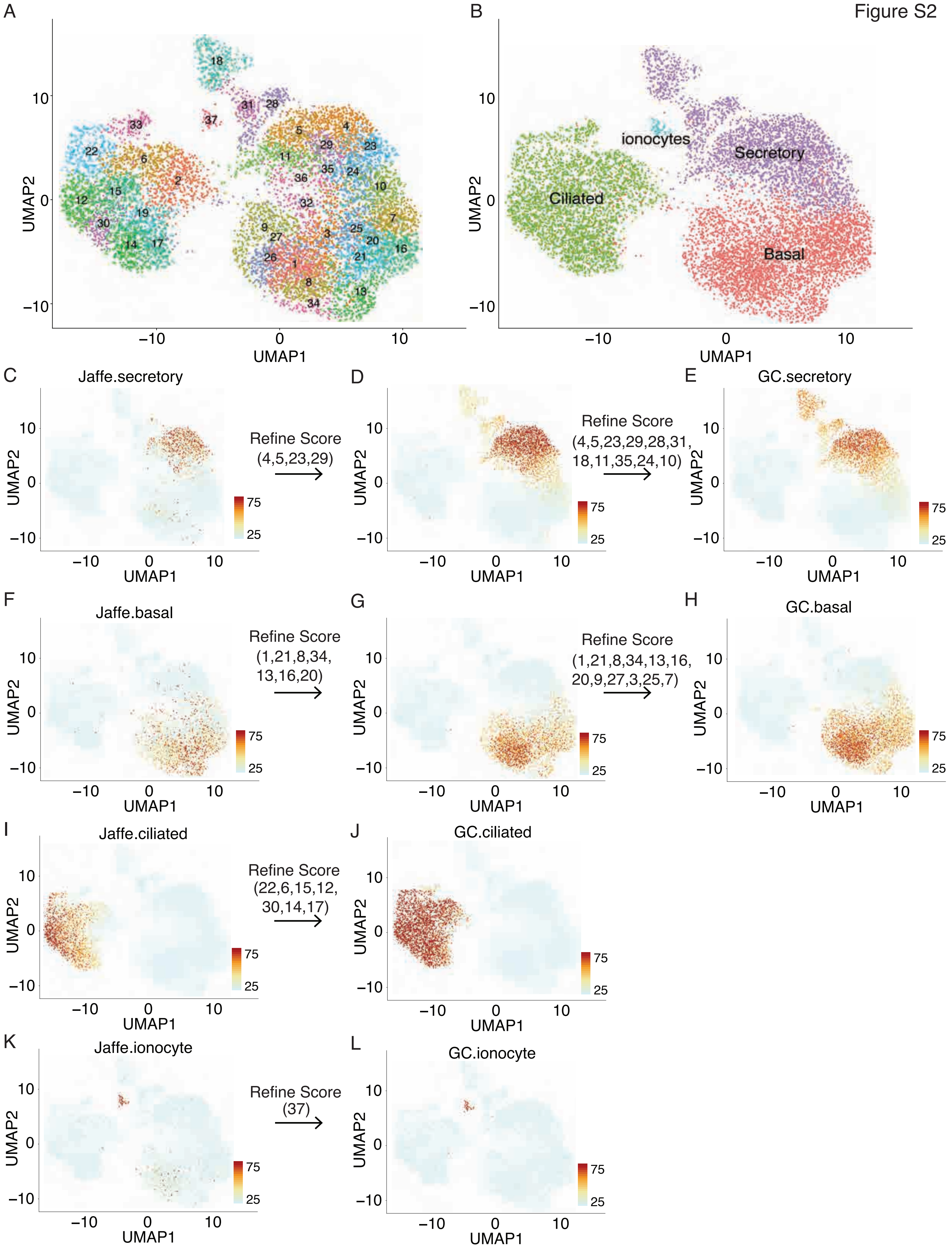
Figure S2
A
B
UMAP1
UMAP2
ionocytes
Secretory
Ciliated
Basal
C
Jaffe.secretory
Refine Score
(4,5,23,29)
D
Refine Score
(4,5,23,29,28,31,
18,11,35,24,10)
E
GC.secretory
F
Jaffe.basal
Refine Score
(1,21,8,34,
13,16,20)
G
Refine Score
(1,21,8,34,13,16,
20,9,27,3,25,7)
H
GC.basal
I
Jaffe.ciliated
Refine Score
(22,6,15,12,
30,14,17)
J
GC.ciliated
K
Jaffe.ionocyte
Refine Score
(37)
L
GC.ionocyte

Figure S3

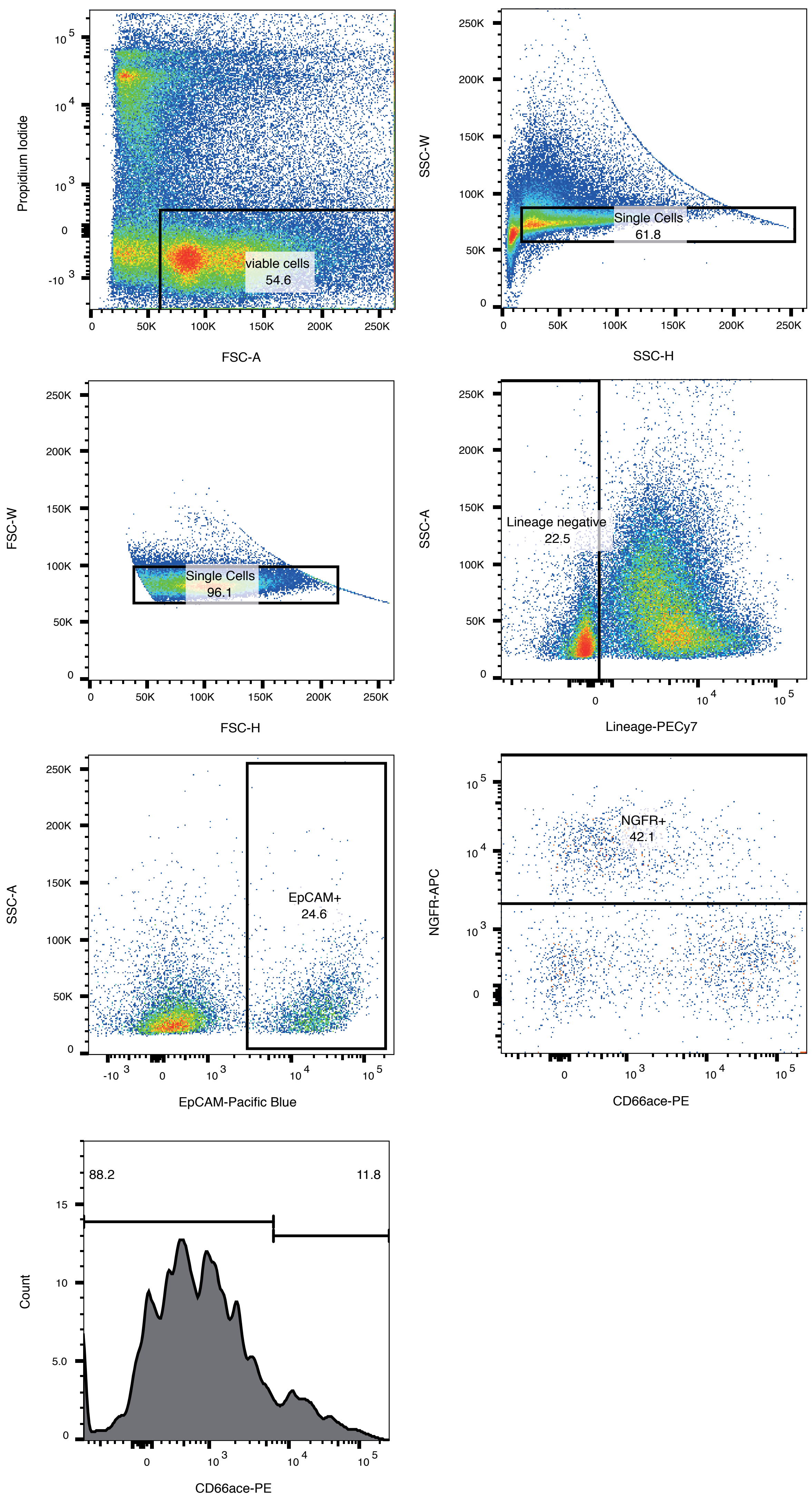

Propidium Iodide
FSC-A
viable cells
54.6
SSC-W
SSC-H
Single Cells
61.8
FSC-W
FSC-H
Single Cells
96.1
SSC-A
Lineage-PECy7
Lineage negative
22.5
SSC-A
EpCAM-Pacific Blue
EpCAM+
24.6
NGFR-APC
CD66ace-PE
NGFR+
42.1
Count
CD66ace-PE
88.2
11.8

Figure S4

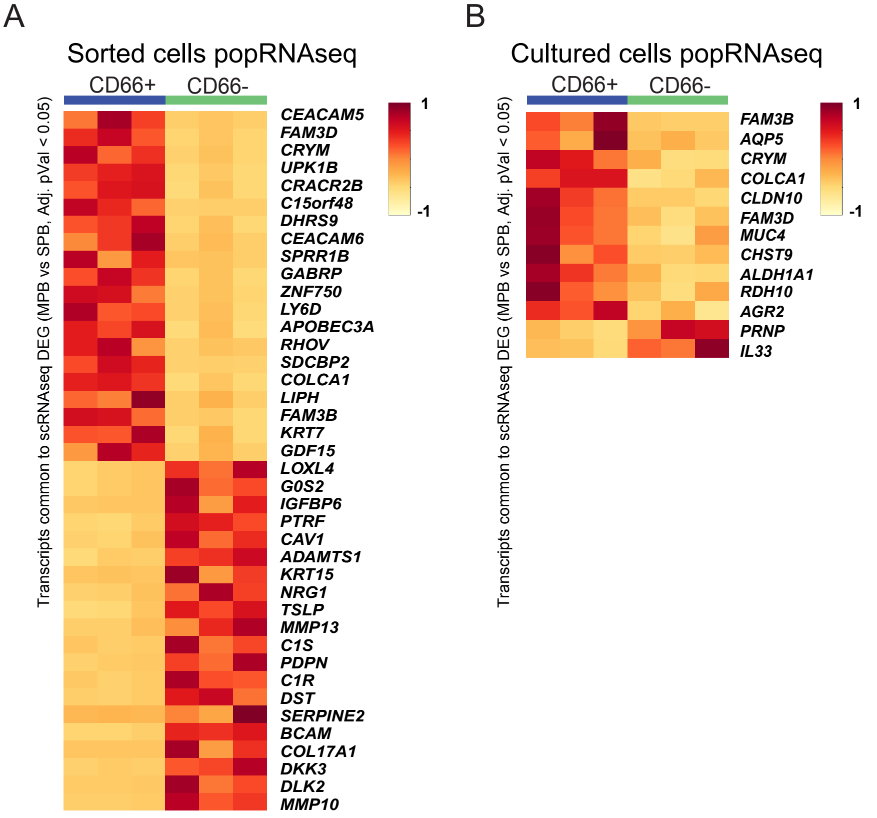
A
Sorted cells popRNAseq
CD66+
CD66-
Transcripts common to scRNAseq DEG (MPB vs SPB, Adj. pVal < 0.05)
CEACAM5
FAM3D
CRYM
UPK1B
CRACR2B
C15orf48
DHRS9
CEACAM6
SPRR1B
GABRP
ZNF750
LY6D
APOBEC3A
RHOV
SDCBP2
COLCA1
LIPH
FAM3B
KRT7
GDF15
LOXL4
G0S2
IGFBP6
PTRF
CAV1
ADAMTS1
KRT15
NRG1
TSLP
MMP13
C1S
PDPN
C1R
DST
SERPINE2
BCAM
COL17A1
DKK3
DLK2
MMP10
1
-1
B
Cultured cells popRNAseq
CD66+
CD66-
Transcripts common to scRNAseq DEG (MPB vs SPB, Adj. pVal < 0.05)
FAM3B
AQP5
CRYM
COLCA1
CLDN10
FAM3D
MUC4
CHST9
ALDH1A1
RDH10
AGR2
PRNP
IL33
1
-1

C

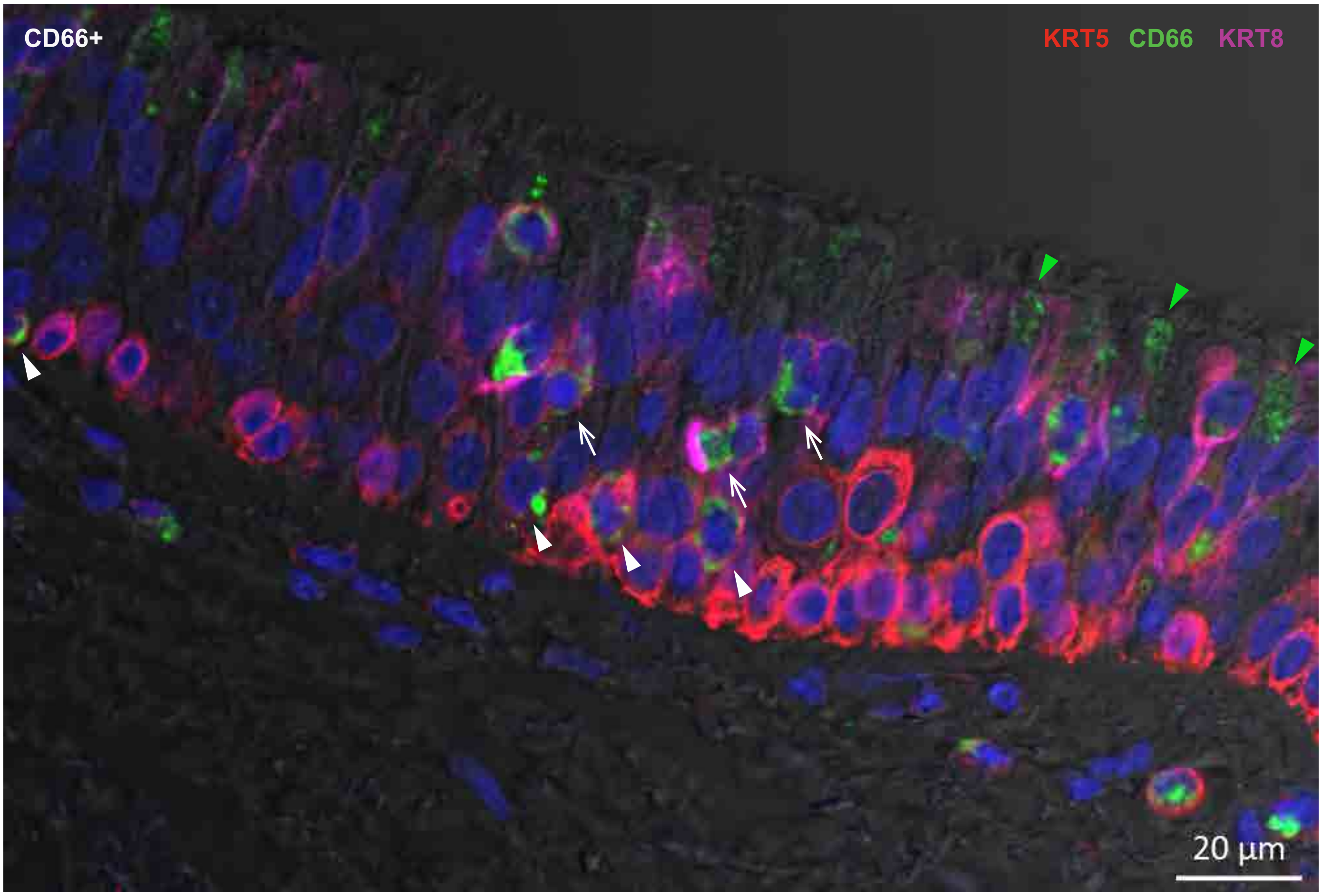
CD66+
KRT5
CD66
KRT8
20 µm

Figure S5

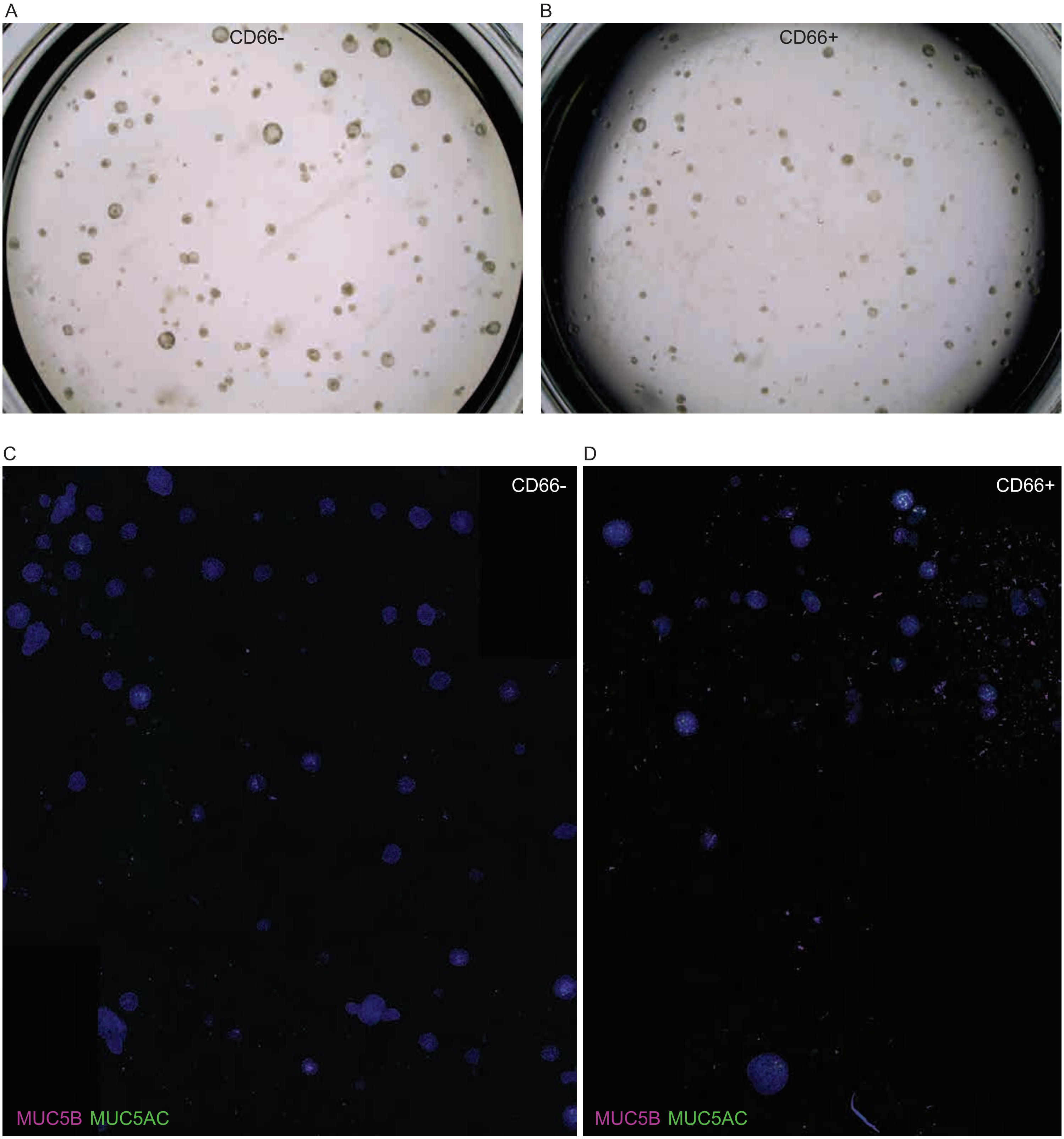
